# A failed top-down control from prefrontal cortex to amygdala in generalized anxiety disorder: evidence from resting-state fMRI with Granger causality analysis

**DOI:** 10.1101/573634

**Authors:** Mengshi Dong, Likun Xia, Min Lu, Chao Li, Ke Xu, Lina Zhang

## Abstract

**Objective:** In generalized anxiety disorder (GAD), abnormal top-down control from prefrontal cortex (PFC) to amygdala is a widely accepted hypothesis through which “emotional dysregulation model” may be explained. However, whether and how the PFC directly exerted abnormal top-down control on amygdala remained largely unknown. We aim to investigate the amygdala-based effective connectivity by using Granger causality analysis (GCA).

**Methods:** Thirty-five drug-naive patients with GAD and thirty-six healthy controls (HC) underwent resting-state functional MR imaging. We used seed-based Granger causality analysis to examine the effective connectivity between the bilateral amygdala and the whole brain. The amygdala-based effective connectivity was compared between the two groups.

**Results:** In HC, the left middle frontal gyrus exerted inhibitory influence on the right amygdala, while in GAD group, this influence was disrupted (single voxel P < 0.001, Gaussian random field corrected with P < 0.01).

**Conclusion:** Our finding might provide new insight into the “insufficient top-down control” hypothesis in GAD.

## Introduction

As the most prevalent anxiety disorder, generalized anxiety disorder (GAD) affects a significant number of people with estimated lifetime prevalence rates of 4-6% (1, 2). The core symptom of GAD is uncontrollable, pervasive worry and anxiety about a variety of events and situations, occurring alongside physical and emotional symptoms, such as restlessness, irritability, muscle tension, difficulties in concentration, sleep disturbance. Although GAD causes massive economic burden and huge individual impacts, the neurobiological basis underpinning the core symptom is poorly understood.

Aberrant amygdala activity and decreased amygdala-prefrontal connectivity are the most consistent neuroimaging findings in GAD, with the latter was considered as the underlying neuropathological mechanism of the core symptom of GAD (3, 4). Decreased functional coupling between amygdala and prefrontal cortex (PFC) is reported in both adults (5, 6) and adolescents with GAD (7, 8). Decreased amygdala-prefrontal coupling also correlated with high anxiety ratings scores (5, 6). Conversely, what’s more interesting is that increased amygdala-prefrontal coupling can predict effective emotion regulation (9) or positive reappraisal of negative emotional material (10).

The hypothesis of insufficient top-down emotion control from PFC to amygdala was proposed (3, 11), when summarized the above findings and given the fact that amygdala is the core hub for emotional processing (12) and PFC is one of the most important regions of the goal-directed attention system (13, 14). However, a contrary hypothesis was also proposed by researchers that GAD involves a hyper-active top-down control system (3, 11). Some studies supporting the second found greater PFC activation during motional regulation process (15, 16). Notably, the two competing hypothesis are based on evidences that altered activation or functional connectivity (Pearson’s correlation coefficient between two BOLD time series) in amygdala or PFC. However, there has previously been little direct evidence that PFC exerts abnormal top-down control on amygdala.

Effective connectivity using Granger causality analysis (GCA) is a very suitable method to investigate the top-down control (directional influence) of PFC on amygdala, because of it can measure the influence of activity in a neural system over that in another (17). In contrast to other methods to measure the effective connectivity, GCA does not require any prior knowledge to select interaction region in advance (17-19). Therefore, it can avoid the model assumption bias. Amygdala-based effective connectivity has been already successfully applied to the social anxiety disorder (20). Between-independent component effective connectivity has also been used to investigate amygdala (19). However, to our knowledge, amygdala-based voxel-wise effective connectivity has not been applied to test the two hypotheses in GAD.

In this article, we aim to investigate the amygdala-based effective connectivity by using GCA. We hypothesize that PFC either exerts hypo-active or hyper-active top-down control on amygdala.

## Material and Methods

### Participants

The study was approved by the Research Ethics Committee of People’s Hospital of Yuxi City (Yuxi, People’s Republic of China). All patients with GAD were recruited from the Department of Rehabilitation Medicine at People’s Hospital of Yuxi City. All participants signed written informed consent agreement. The inclusion criteria for GAD patients were: (a) must conform to Diagnostic and Statistical Manual of Mental Disorders, Fourth Edition (DSM-IV) for diagnosis of GAD; (b) complaining of excessive, uncontrollable and lasting anxiety and worry about everyday events for at least 6 months; (c) free of any anti-anxiety or anti-psychotic drugs at least 1 month before and during the study; (d) had a Hamilton Anxiety Rating Scale (HAM-A) score ≥ 14; (e) younger than 60 years old and (f) right-handedness. Exclusion criteria for GAD patients were: (a) had focal abnormality in the structural MRI; (b) had other psychiatric disorders or secondary anxiety disorder or substance abuse; (c) a past history of depression or episodes of depression; (d) diagnosed with other serious disease, such as diabetes or high blood pressure; (e) being pregnant or breastfeeding and (f) head motion more than or equal to 2.0 mm or 2.0°during MR imaging. Four GAD patients were excluded because with abnormal signals in the structural MRI. Four GAD patients were excluded because head motion. Finally, 35 GAD patients were included in the study.

The study also included 36 gender-, age-, and hand-matched healthy controls (HC). The inclusion criteria for healthy controls were as follows: (a) no history of neurological or psychiatric disorder; (b) no focal abnormality in the structural MRI; (c) had a HAM-A score < 7 and a Hamilton Depression Rating Scale (HAM-D) score < 8 and (d) head motion less than 2.0 mm or 2.0°during MR imaging.

### Clinical Measures

Before fMRI scan, all participants completed HAM-A, the HAM-D and State–Trait Anxiety Inventory (STAI) scales.

### Data Acquisition

Scanning took place on a 3.0 T MRI scanner with an eight-channel head coil (Ingenia; Philips, the Netherlands) at People’s Hospital of Yuxi City. Foam pads was used to minimize head motion. During data acquisition, all subjects were instructed to keep their eyes closed, relax mind, hold still, and not to fall asleep. The resting-state fMRI were acquired using a gradient echo planar imaging sequence (EPI). The sequence parameters were as follows: repeat time/echo time = 2000 ms/35 ms, field of view =230×230 mm, flip angle = 90°, in-plane matrix =64×64, 35 transverse slices with thickness = 3.6 mm and slice gap = 0.7 mm, voxel size = 3.6×3.6×3.6 mm^3^. For each participant, 240 volumes were acquired, resulting in a total scan time of 480 s.

### Data Preprocessing

Data were preprocessed using the SPM12 and the Data Processing Assistant for Resting-State fMRI software (DPARSF, Advanced Edition, V4.3) (http://www.rfmri.org/DPARSF). The first 10 images of each subject were discarded. Then, the resting-state fMRI data were corrected for the temporal differences between slices and head motion. Those Participants whose head movement exceeded 2.0 mm or 2.0°were excluded. Next, the corrected fMRI data were spatially normalized to the standard Montreal Neurological Institute (MNI) template, and were resampled to 3×3×3 mm^3^. These data were spatially smoothed with a 6 mm full-width at half maximum isotropic Gaussian kernel. We further processed the data to remove linear trends and filtered temporally (band-pass, 0.01–0.1Hz). Finally, nuisance signals including 24 head motion parameters, CSF signals, white-matter signal, and global signal were regressed out from the fMRI data.

### Granger Causality Analysis

The bilateral amygdala in the automated anatomical labeling (AAL) template were used as the seed regions for the Granger causality analysis. Bivariate first-order coefficient-based voxel-wise GCA was implemented on the REST software (http://www.restfmri.net/forum/REST-GCA), using signed-path coefficients (21). We followed Chen’s extended model as following:

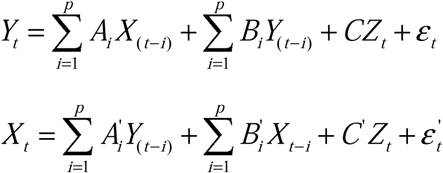

where: *Y*_*t*_ is the BOLD time series of one voxel in the brain at time *t*; *Xt* is the BOLD time series of seed region at time *t*; *Z*_*t*_ is a *q*×1 vector containing exogenous variables (covariates or confounds) at time *t*; *ε*_*t*_ is the error term; *p* and *q* are the number of lags and confounds respectively; *A*_*i*_ is the signed path coefficient at time lag *i* (*i*=1,…, *p*); *B*_*i*_ is the autoregression coefficient. In the present study, the number of lags *p* = 1 (1 TR = 2 s).

If *A*_*i*_ is significantly different from zero, then it is said that *X* Granger causes *Y*. If The path coefficient *A*_*i =*_ +1, it suggests that one unit of change in the time series *X* in a specific direction brings a unit change in the time series Y in the same direction, representing excitatory influence and vise verse (22). It is worth noting that the voxel-wise GCA calculates causal influences of a time series of the seed region on that of every voxel in the brain (X to Y) and the time series of every voxel in the brain on that of the seed region (Y to X). Thus, 4 GCA maps (2 seed regions ×2 directions of GCA influences) were generated for each subject.

### Statistical Analysis

Difference in age between GAD and HC group was compared by using Mann-Whitney U-test. Between-group differences in STAI, HAM-A and HAM-D were compared by using two-sample t-test. Difference in gender was assessed by using chi-squared test.

Firstly, the 4 GCA maps were analyzed using one-sample t test for GAD and HC group separately with an uncorrected P < 0.001 and cluster size > 10. Then, we made 4 masks by combining every 2 resulted maps (GAD + HC) from the above one-sample t test to constrain the subsequent two-sample t-test.

Secondly, Between-group differences in GCA maps were compared by using two-sample t test with age and gender imported as covariates. Multiple comparisons were corrected by a Gaussian random field (GRF) method implemented in the DPABI software (DPABI version 3.0, Data Processing & Analysis for (Resting-State) Brain Imaging) (23) and using significant corrected thresholds of P < 0.01 with combined with single voxel P < 0.001.

For regions showing significant group differences, mean signed-path coefficients were extracted and follow-up one-sample t tests were performed to investigate the Granger causal influences in each group separately.

## Results

### Demographic and Clinical Variables

As Table 1 shown, the GAD patients and HC showed no significant differences in gender (*P* = 0.93) and age (*P* = 0.67). GAD group had higher STAI, HAM-A and HAM-D than those of HC group (all *P* < 0.001).

**Table 1.** Demographic, clinical and neuropsychological characteristics of the participants

| **Characteristics** | **GAD**<br>**(n=35)** | **HC**<br>**(n=36)** | ***P* Value** |
| --- | --- | --- | --- |
| Gender (M/F) | 12/23 | 12/24 | 0.93<sup>a</sup> |
| Age (y) | 39.43 ± 10.61 | 38.69 ± 9.20 | 0.67<sup>b</sup> |
| Duration (mo) | 29.69 ± 43.86 | N/A | N/A |
| STAI |  |  |  |
| State | 51.34 ± 11.59 | 21.17 ± 2.48 | < 0.001<sup>c</sup> |
| Trait | 50.14 ± 9.85 | 21.19 ± 2.81 | < 0.001<sup>c</sup> |
| HAM-A | 22.69 ± 6.21 | 1.00 ± 1.80 | < 0.001<sup>c</sup> |
| HAM-D | 16.69 ± 6.05 | 0.39 ± 1.23 | < 0.001<sup>c</sup> |

Note.—Unless otherwise noted, data are mean ± standard deviation.
GAD, Generalized Anxiety Disorder; HC, Healthy Controls; STAI, State-trait Anxiety Inventory; HAM-A, Hamilton Anxiety Rating Scale; HAM-D, Hamilton Depression Rating Scale; N/A, Not Available.
<sup>a</sup> The *P* value was obtained by using chi-square test. <sup>b</sup> The *P* value was obtained by using Mann-Whitney U-test. <sup>c</sup> The *P* value was obtained by using two-sample t-test.

### Granger Causality Analysis

Compared with HC, GAD patients showed disrupted Granger causal influences from the left middle frontal gyrus (dorsolateral prefrontal cortex, DLPFC) to the right amygdala (Table 2 and Figure 1). Follow-up one-sample t tests found that, in controls, the left DLPFC exerted a significant inhibitory influence on right amygdala (P < 0.001), while in the GAD patients, this influence was not statistically significant (P = 0.25) (Figure 2).

**Table 2.** Between-Group Differences in Effective Connectivity

| Brain Regions | MNI Coordinates | Cluster Size<br>(Voxels) | *T* Values<br>(peak) | Mean Causal Influences |  |
| --- | --- | --- | --- | --- | --- |
|  | (x, y, z) |  |  | HC | GAD |
| MFG.L→AMYG.R | (-36 45 27) | 31 | 4.63 | -1.14±1.24<br>p<0.001<sup>#</sup> | 0.21±1.09,<br>p=0.25<sup>#</sup> |

GAD, Generalized Anxiety Disorder; HC, Healthy Controls;
<sup>#</sup>, one-sample t test

**Figure 1:**
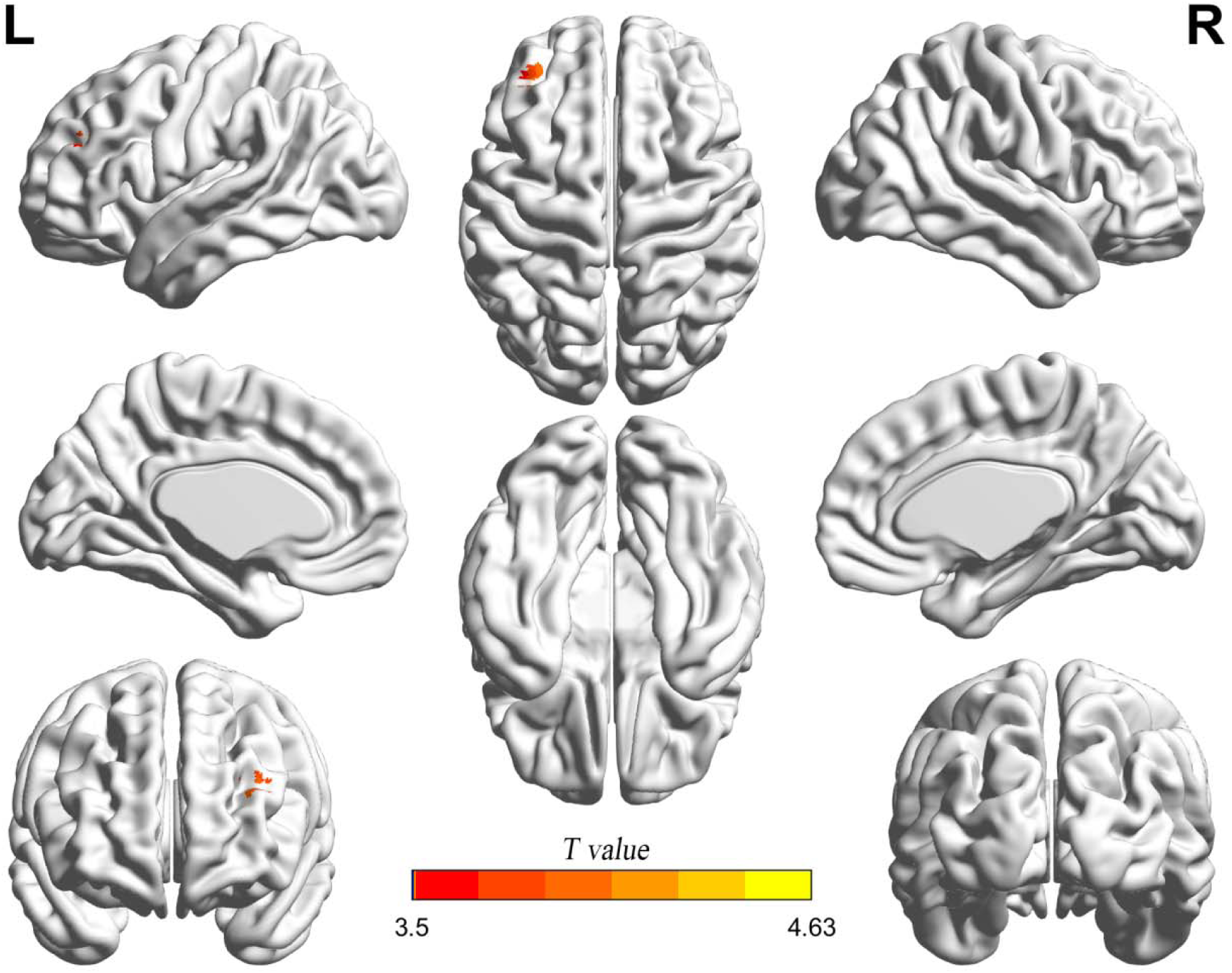
Between-Group Differences in Effective Connectivity. Compared with HC, patients with GAD showed abnormal Granger causal influences from the left middle frontal gyrus to the right amygdala.

**Figure 2:**
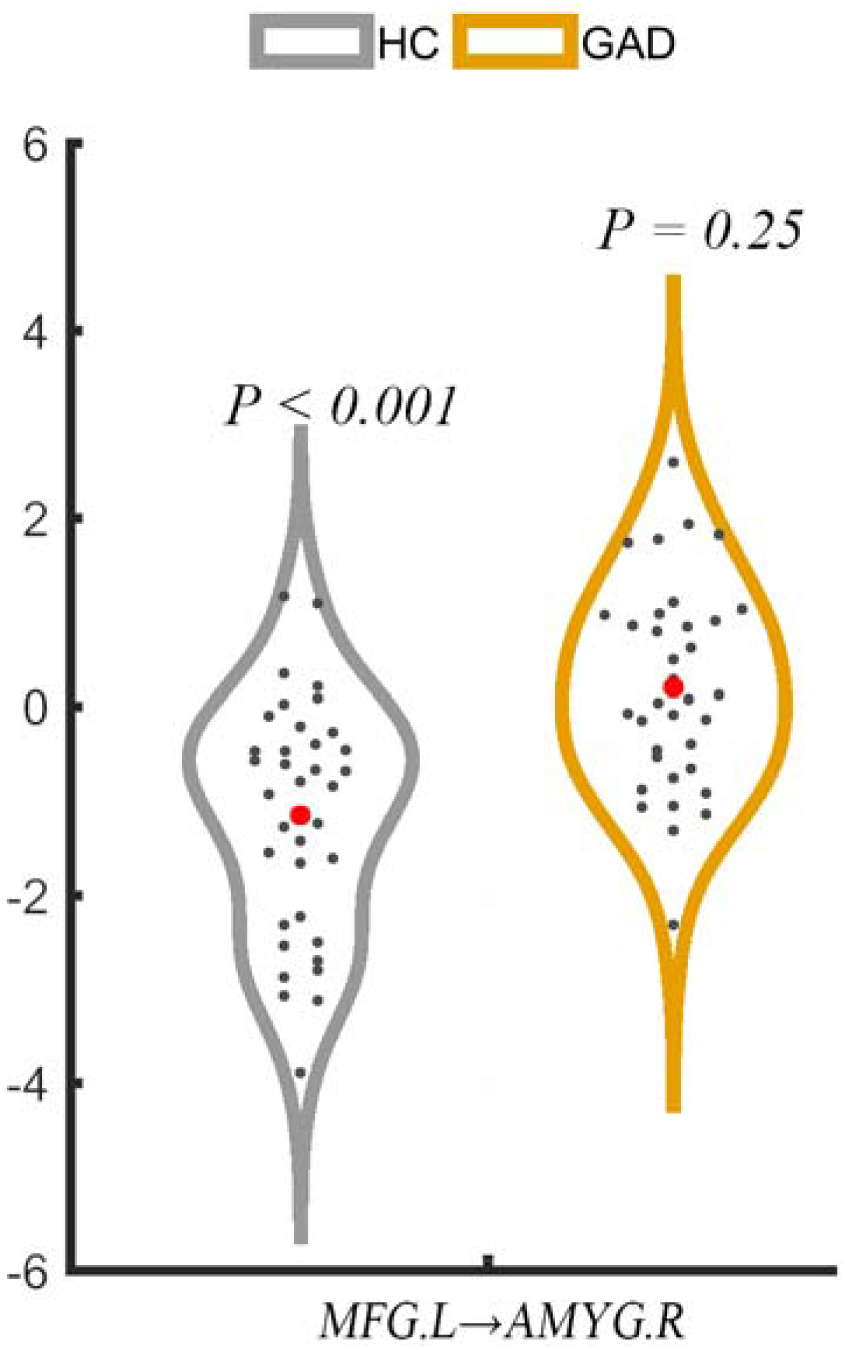
Follow-up one-sample t tests found that, in controls, the left middle frontal gyrus exerted a significant inhibitory influence on the right amygdala (P < 0.001), while in the GAD patients, this influence was not statistically significant (P = 0.25). MFG.L, Left middle frontal gyrus; AMYG.R, right amygdala.

## Discussion

This study investigated amygdala-based effective connectivity (directional connectivity) in HC and patients with GAD at resting state. Our main finding showed that, in HC, the left DLPFC exerted a significant inhibitory influence on right amygdala (significant negative coefficient), however, in patients, the inhibitory influence was disappeared (no significant coefficient). This suggests a failed top-down control from DLPFC to amygdala in GAD at resting state. Our results align with the “insufficient top-down control” hypothesis and provide some new insight into the pathophysiological mechanism of GAD.

In line with “insufficient top-down control” hypothesis, many studies found decreased activation in PFC or decreased functional connectivity between PFC and amygdala (3-8, 11). A recent study using functional network modeling and graph-theory found lower betweenness centrality and lower degree in PFC, also suggesting weakened inhibitory control of PFC. Besides, using average time courses in independent component, Jianping Qiao et al. found the effective connectivity in GAD was decreased from middle frontal gyrus to amygdala (19). In the present study, using effective connectivity with voxel-wise analysis, we also found a failed top-down control from left DLPFC to right amygdala in GAD with relatively large sample. Together with the finding by Jianping Qiao et al. (19), our finding might deepen the understanding of the “insufficient top-down control” hypothesis.

Decreased functional coupling between PFC and amygdala is a reasonable explanation for the two common abnormalities in GAD: decreased activation of PFC, increased activation of amygdala (3, 4, 6, 24, 25). Disrupted top-down control from DLPFC to amygdala might assist the decreased functional coupling to explain these two consistent findings in GAD. In HC, the DLPFC exerts an inhibition influence on amygdala. If the causal link between them is broken in GAD, the amygdala is very possible to lose control and showed increased activation. On the contrary, the prefrontal cortex might lose positive feedback, thereby reducing activation. Future studies will be needed to test this hypothesis.

Our study had several limitations. First, this study is a cross-section study, and the findings cannot provide longitudinal alteration data on patients with GAD. Second, we cannot clearly identify the effective connectivity during emotion regulation tasks due to that the present study is resting-state fMRI study.

Overall, our data offers an explanation for the “insufficient top-down control” hypothesis in GAD.

